# Loops linking secondary structure elements affect the stability of molten globule intermediate state of apomyoglobin

**DOI:** 10.1101/857102

**Authors:** M.A. Majorina, B.S. Melnik

## Abstract

Apomyoglobin is a protein widely used as a model for studying globular protein folding. This work aimed to test the hypothesis on influence of rigidity and length of loops linking protein secondary structure elements on the stability of molten globule intermediate state. For this purpose, we studied folding/unfolding of mutant apomyoglobin forms with substitutions of proline residues to glycine and with loops elongated by three and six glycine residues. For all the protein forms, denaturation/renaturation kinetic curves at different urea concentrations were obtained, folding/unfolding constants were calculated and dependencies of rate constant logarithms on urea concentrations were plotted. All the data gave an opportunity to calculate free energies of different apomyoglobin states. As a result, the mutations in apomyoglobin loops were demonstrated to have a real effect on intermediate state stability compared to unfolded state.

## Introduction

Apomyoglobin is a convenient model for studying globular protein folding/unfolding *in vitro*. This small alpha-helical of 17 kDa molecular mass consists of 153 amino acid residues, of which 65% form eight α-helices [1]. The structure of this protein is shown on Fig. 1. So far, apomyoglobin has been studied quite well. Numerous studies allowed determination of a number of its basic characteristic features. Particularly, the process of apomyoglobin folding is divided into two stages, fast and slow, the first one being related to very fast formation of molten globule intermediate state, possessing a hydrophobic core with native-like helical structure [2, 3]. The slow phase is related to protein transition from molten globule into native state [2]. Kinetic experiments based on measurement of stopped-flow tryptophan fluorescence do not let calculate molten globule formation rate (i.e. rate constants of rapid stage of folding), because it is formed in «dead time» of the instrument, around several microseconds. An experimental approach for studying kinetics of multistage protein folding was developed in our laboratory, allowing to obtain data on occupancy of the intermediate state of the protein as a function of denaturating agent concentration. The data obtained by means of such an approach could be used for estimation of the stability of molten globule state.

**Figure 1.**
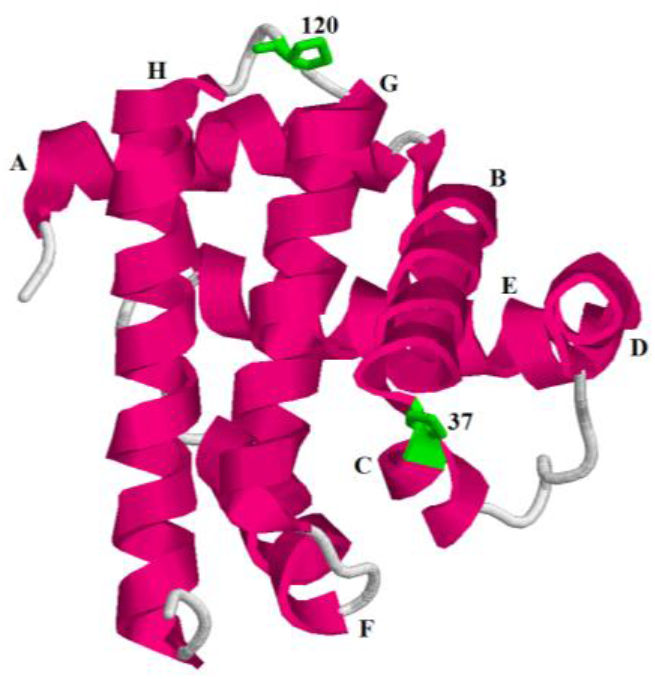
3D structure of myoglobin (heme not shown) (PDB ID: 1BZP). Polypeptide chain trace is shown in ribbon mode. Eight alpha-helices are designated by letters from A to H. Side chains of proline residues P37 and P120 are shown in wires.

It is important to note that the structure of kinetic intermediate of apomyoglobin folding is analogous to the molten globule observed at pH 4.2 in equilibrium experiments [3].

Moreover, NMR studies show that some mutations are capable of changing the direction of apomyoglobin folding [4], because they can alter the structure of kinetic intermediate of folding [5].

In previous works of our laboratory, systematic investigations were conducted in order to reveal the effect of various mutations on apomyoglobin energy profile [2, 6–9]. Circular dichroism and fluorescence measurement showed that none of 20 studied single amino acid substitutions in the core or on the myoglobin surface did alter the stability of molten globule intermediate state [6, 8, 9]. It should be highlighted that the similar intermediate state appears during folding of the majority of the proteins. Thus, knowledge of its stability and folding features is necessary to understand the self-assembly of most of the protein molecules.

*There appears a question: is it possible to alter the stability of apomyoglobin intermediate state by any mutations?*

We supposed that mutations affecting the mobility of the main chain and, consequently, mobility and compactness of the spatial structure of protein intermediate states could have such effect, for example, amino acid substitutions in loops linking myoglobin secondary structure elements.

In this study we investigated the effect of insertion of glycines increasing the flexibility of loops and elongation of the loops by several glycine residues. Such mutations could alter the interaction of secondary structure elements in molten globule state.

To test this hypothesis, we designed the following mutations in apomyoglobin: proline substitution to glycine in two loops (P37G, P120G) and loop elongation in 120 amino acid region by 3 and 6 glycine residues (P120(3G), P120(6G)). The loop including P120 is situated between helices G and H (Fig. 1) formed during myoglobin transition from unfolded state to molten globule state [3]. The loop including P37 connects helices B and C, which are formed during transition from molten globule state to the native state [3].

## Experimental part

Based on a plasmid containing a wild-type apomyoglobin gene, plasmids containing mutant apomyoglobin gene variants with selected substitutions were derived. Recombinant proteins were isolated and purified according to methods described in our earlier works [2, 6–9]. The effect of mutations on the structure of the native state of apomyoglobin was estimated by means of circular dichroism (CD) spectra in far ultraviolet (UV) area. Far UV CD spectra of the proteins were registered with a J-600 spectropolarimeter (Jasco, Japan) at a protein concentration of 1 mg/mL using a 0.1-mm pathway quartz cell. Slight differences between CD spectra of mutants and wild-type protein evidence that the introduced amino acid substitutions had literally no effect on the structure of apomyoglobin (spectra not shown).

Kinetic measurements were taken using a spectrofluorimeter Chirascan Spectrometer (Applied Photophysic, UK) equipped with a stopped-flow attachment. The excitation wavelength was 280 nm, and emission spectra were recorded using a 320-nm cut-off glass optical filter. The initial urea concentration was 5.5 M for refolding experiments, and 0.0 M for unfolding ones. The initial protein solution was mixed (1:1) with a buffer of various urea concentrations using the stopped-flow attachment. The final protein concentration was 0.03 mg/mL. All the measurements were carried out at 11°C in 20 mM sodium phosphate buffer, pH = 6.2. Denaturation/renaturation kinetic curves at different urea concentrations (0-5.5M) were obtained for each protein variant. The technique of the measurements was described in detail in our previous works [2, 6–9].

## Results and discussion

One should keep in mind that apomyoglobin folding includes fast and slow stages. The fast folding phase corresponds to the transition of the protein from unfolded state to intermediate state (U↔I). This phase takes place in “dead time” of the instrument, thus it is impossible to measure it. The experimentally derived folding curves of apomyoglobin allow calculation of only slow folding phase rate constant, which corresponds to protein transition from intermediate state (I) to native state (N). To calculate apparent rate constants, kinetic curves of folding/unfolding slow phase were approximated in Sigma Plot program package according to formula (1),

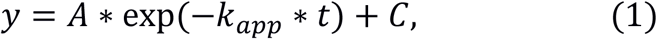

where *k*_*app*_ is the value of folding/unfolding apparent rate constant; *A* and *C* are curve parameters; *t* is reaction time.

After calculation of apparent rate constants, chevron plots (dependencies of logarithms of apparent rate constants on denaturant concentrations) can be made.

Figure 2A-E shows chevron plots for wild-type apomyoglobin and its four mutant variants.

**Figure 2.**
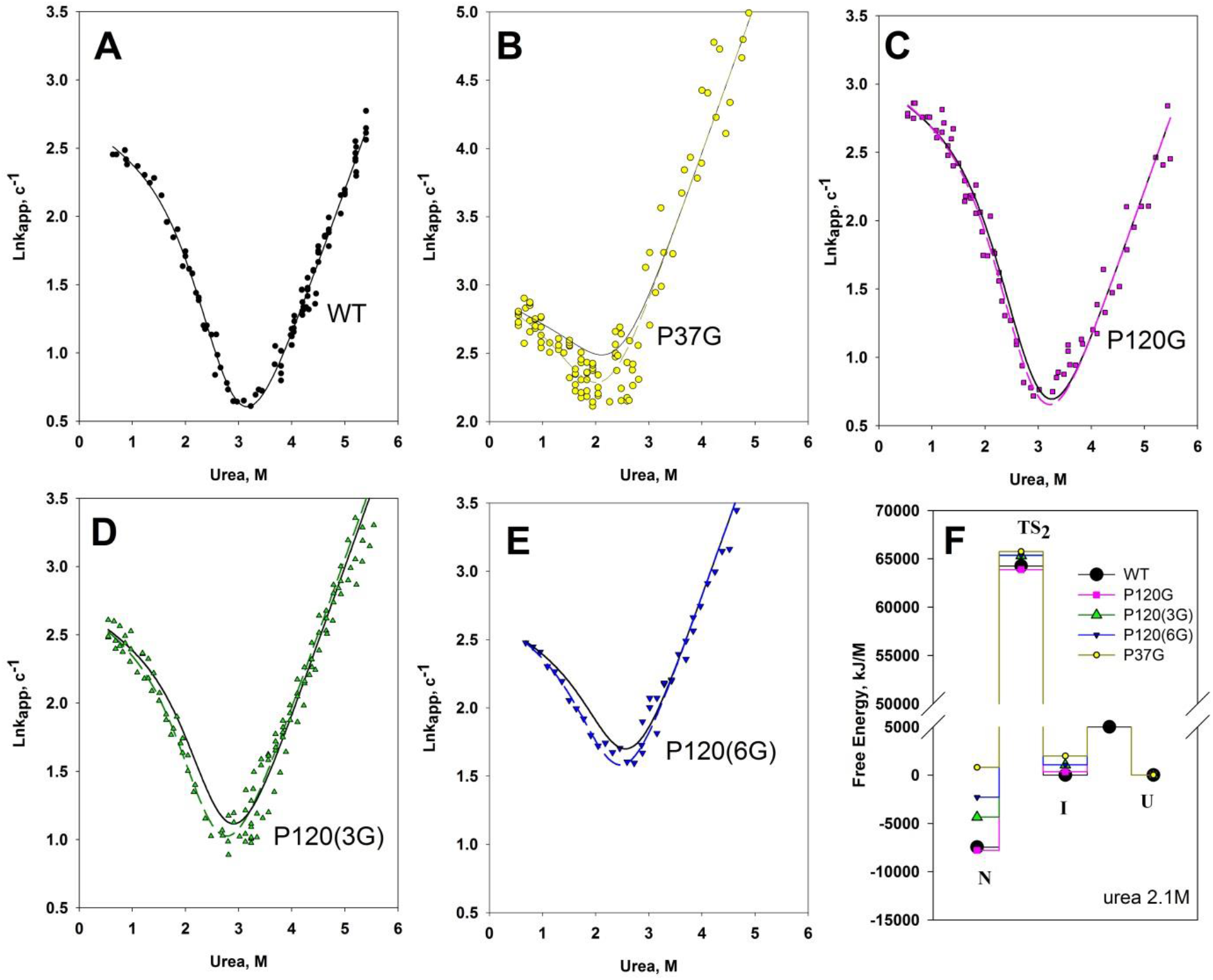
A,B,C,D,E — chevron plots of dependencies of apparent folding/unfolding rate constant of wild-type apomyoglobin (WT) and studied mutant forms (legends on the plot) on denaturant concentration. F - free energy profiles built from approximation of chevron plots.

Each point of chevron plots corresponds to natural logarithm of apparent rate constant of folding/unfolding calculated by formula (1) at given urea concentration.

For proteins with one-stage folding model, the apparent rate constant on chevron plots equals the sum of forward and reverse reaction rate constants:

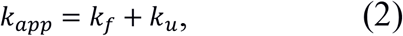

where *k*_*app*_ – is folding/unfolding apparent rate constant, *k*_*f*_ is folding rate constant, *k*_*u*_ is unfolding rate constant.

In the case of myoglobin, which has two-stage folding model, an additional parameter *F*_*I*_, occupancy of intermediate state [2], must be introduced into the formula (2) to take into account the influence of fast folding phase onto the subsequent slow phase:

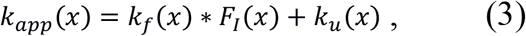

where *F*_*I*_ (*x*) is occupancy of the intermediate state (value in range from 0 to 1); *x* is urea concentration.

The equation used for chevron plot approximation can be written in general form:

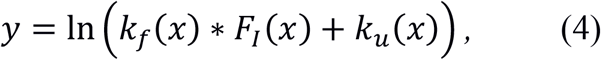

where

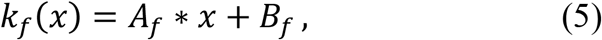

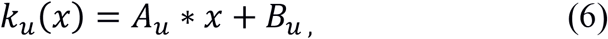

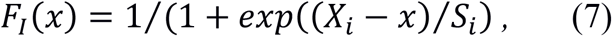

*A*_*f*_, *B*_*f*_, *A*_*u*_, *B*_*u*_ are the parameters of linear functions depicting real rate constants of slow phase of apomyoglobin folding/unfolding, *X*_*i*_ and *S*_*i*_ are parameters of S-shaped curves depicting occupancy of the intermediate state; *X*_*i*_ is the center of the sigmoid, *S*_*i*_ — slope of sigmoid [2].

If supposed that the track of folding of mutant apomyoglobin variants did not alter compared to the wild-type protein, then every chevron plot must have the same slope of folding branches (*A*_*f*_), the same slope of unfolding branches (*A*_*u*_) and the same occupancy parameters for intermediate state, *X*_*i*_ and *S*_*i*_ [2]. Only the position of unfolding and folding brances, i.e. parameters *B*_*f*_ and *B*_*u*_, should change, but they can be quite accurately determined experimentally at low and high urea concentrations (“weights” of the points at the borders of the plot are higher than in the middle). Thus, we use quite a few restrictions during approximation of chevron plots. In other words, during approximation of data for mutant proteins, all the parameters in equations 4-7 are “fixed”.

Solid black line on Figure 2 shows the approximation of experimental data (chevron plots) with the aforementioned restrictions in the equation, i.e. with restrictions supposing that the mutations did not alter the stability of apomyoglobin intermediate state. It could be seen that the middle part of chevron plots is poorly described for mutant proteins. The greatest differences are observed for the protein with substitution of proline 37 to glycine (Fig. 1B) and for the protein with a loop elongated by six glycines in position 120 (Fig. 1E). Such differences could mean that the studied mutations altered exactly the stability of intermediate apomyoglobin state compared to unfolded state.

When a restriction on just one parameter related to the stability of intermediate state, i.e. *X*_*i*_, is “removed”, all the experimental data are approximated very well (dashed line on Figure 2). Thus, one can conclude that the studied mutations did alter the stability of intermediate state of apomyoglobin compared to its unfolded state.

Approximation of chevron plots allows calculating the parameters (*k*_*u*_, *k*_*f*_, *F*_*I*_), which, in turn, allow building the free energy profiles clearly showing what exact protein state was altered by each mutation (Fig. 2, 3)

**Figure 3.**
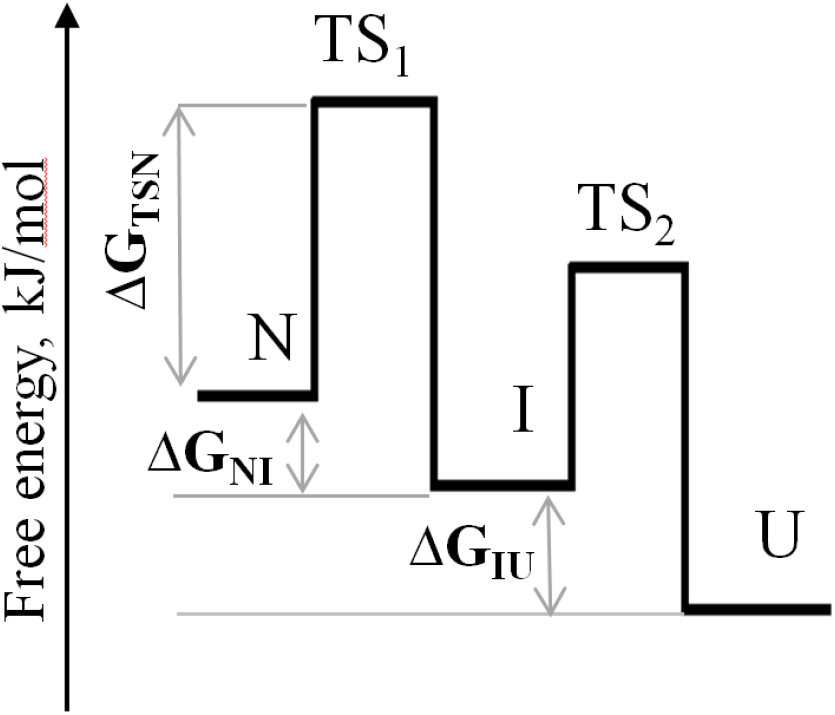
A scheme of free energy plot (N is the native state of the protein, I is the intermediate state, U is the unfolded state, TS_1_ and TS_2_ are the first and the second transition states, ΔG_TSN_, ΔG_NI_, ΔG_IU_ are changes in free energy)

Formulas for building free energy profiles were as follows:

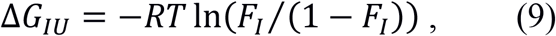

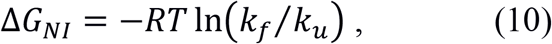

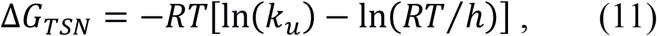

where *F*_*I*_ is the occupancy of intermediate state; *k*_*f*_ is unfolding rate constant, *R* is universal gas constant, *T* is temperature expressed in К, *h* is Planck constant.

Figure 2F shows free energy profiles of wild-type apomyoglobin and studied mutant forms in 2.1 M urea. This is the concentration at which free energies of unfolded state and molten globule intermediate state are equal in wild-type protein.

One can see that the studied mutations of apomyoglobin affect the stability of native (N) and transition state (TS2) of the protein in different maner, but all of them more or less destabilize the intermediate state (MG) compared to unfolded (U) protein state.

## Main results and colclusions

We studied folding/unfolding of mutant apomyoglobin forms with proline substitutions to glycine in two of its loops (P37G and P120G), as well as with elongation of the loop in position 120 by three and six glycine residues (P120(3G) and P120(6G)). For all the mutant protein forms, chevron plots (dependencies of apparent rate constants of denaturation/renaturation on denaturant concentration) were made and free energies of native, transition and intermediate states were calculated. The results of the experiments showed that the studied mutations in loops linking apomyoglobin secondary structure elements destabilize molten globule intermediate state compared to the unfolded state.

Thus, the increased flexibility of protein loops affects the stability of molten globule intermediate state.

## Acknowledgement

The work was supported by Russian Foundation for Basic Research, grant number 18-34-00307.

